# A plasma membrane-associated glycolytic metabolon is functionally coupled to K_ATP_ channels in pancreatic α and β cells from humans and mice

**DOI:** 10.1101/2022.10.06.511200

**Authors:** Thuong Ho, Evgeniy Potapenko, Dawn B. Davis, Matthew J. Merrins

## Abstract

The ATP-sensitive K^+^ channel (K_ATP_) is a key regulator of hormone secretion from pancreatic islet endocrine cells. Using direct measurements of K_ATP_ channel activity in pancreatic β cells and the lesser-studied α cells, from both humans and mice, we demonstrate that a glycolytic metabolon locally controls K_ATP_ channels on the plasma membrane. The two ATP-consuming enzymes of upper glycolysis, glucokinase and phosphofructokinase, generate ADP that activates K_ATP_. Substrate channeling of fructose 1,6-bisphosphate through the enzymes of lower glycolysis fuels pyruvate kinase, which directly consumes the ADP made by phosphofructokinase to raise ATP/ADP and close the channel. We further demonstrate the presence of a plasma membrane NAD^+^/NADH cycle, whereby lactate dehydrogenase is functionally coupled to glyceraldehyde-3-phosphate dehydrogenase. These studies provide direct electrophysiological evidence of a K_ATP_-controlling glycolytic signaling complex and demonstrate its relevance to islet glucose sensing and excitability.

**Graphical abstract:** Ho *et al*. demonstrate that the enzymes of glycolysis, as well as lactate dehydrogenase, form a plasma membrane-associated metabolon with intrinsic ATP/ADP and NAD^+^/NADH cycles. The subcellular location of this signaling complex allows both ATP-consuming and ATP-producing enzymes to locally control the activity of ATP-sensitive K^+^ channels in pancreatic α and β cells from humans and mice.

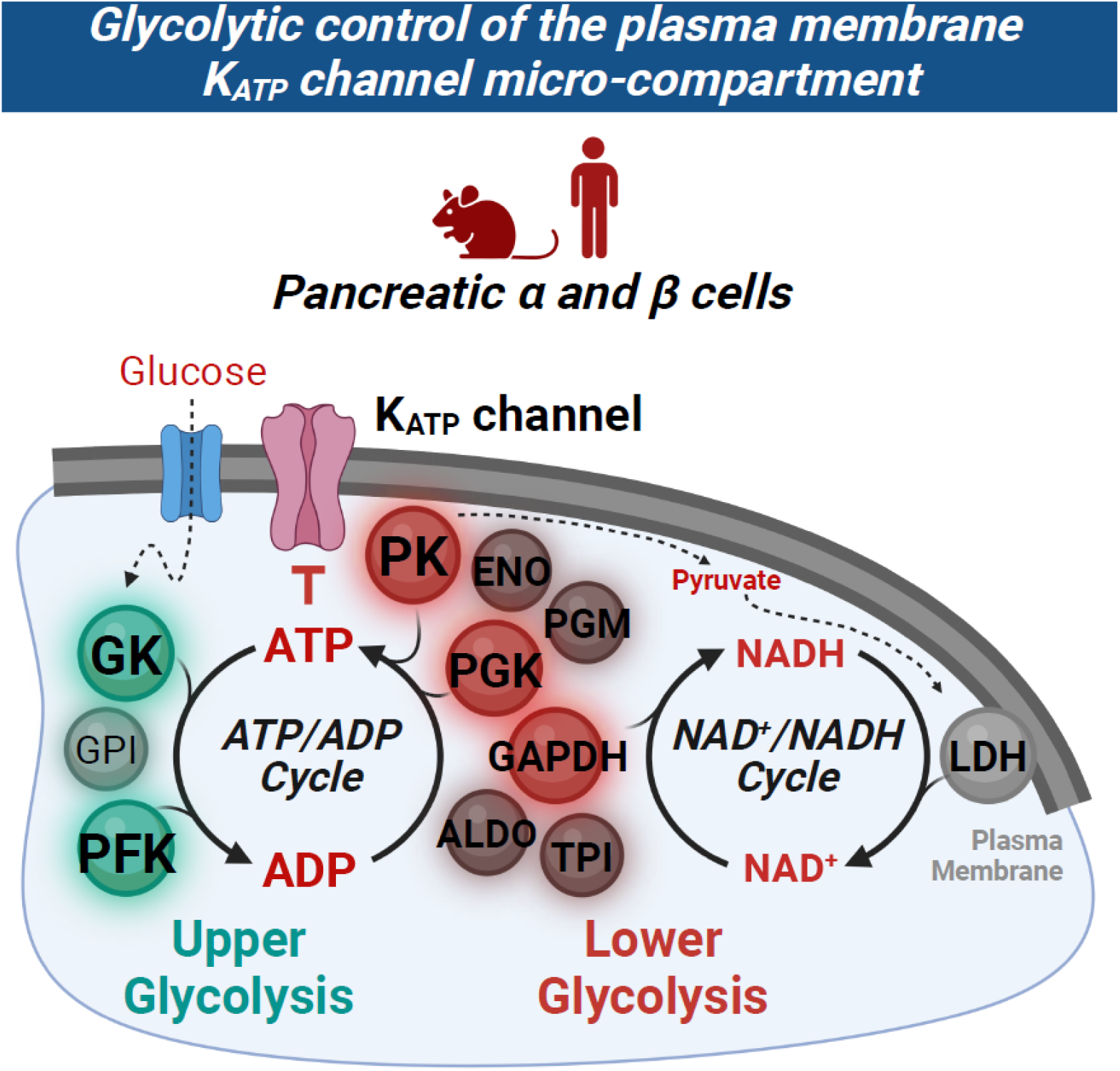

**Highlights:**

- K_ATP_ channels are regulated by a glycolytic metabolon on the plasma membrane.
- Substrate channeling occurs between the consecutive enzymes of glycolysis.
- Upper glycolysis produces ADP that is used directly by lower glycolysis to make ATP.
- LDH and GADPH facilitate a plasma membrane-associated NAD^+^/NADH redox cycle.

## Introduction

The ATP-sensitive K^+^ channel (K_ATP_) has a critical role in controlling insulin and glucagon release from pancreatic islet endocrine cells, which are essential for the regulation of blood glucose homeostasis. K_ATP_ channels are composed of pore-forming Kir6.2 subunits that are inhibited by ATP, and regulatory SUR1 subunits that are activated by MgADP, such that elevation of the sub-plasma membrane ATP/ADP ratio (ATP/ADPp_m_) leads to K_ATP_ channel closure^1–3^. In pancreatic β cells, closure of K_ATP_ channels induces membrane depolarization, resulting in the activation of voltage-sensitive Ca^2+^ channels that trigger insulin exocytosis^4–6^. In pancreatic α cells, K_ATP_ channels exist predominantly in the closed state, while full closure during hyperglycemic conditions inactivates voltage-dependent Na^+^ channels and subsequently suppresses Ca^2+^-stimulated glucagon release^7,8^. Thus, in both cell types, the K_ATP_ channel serves as a metabolic sensor for the plasma membrane that determines cellular excitability.

While it has been historically assumed that K_ATP_ channel closure is mediated by mitochondrial ATP generated via oxidative phosphorylation, recent evidence suggests that β cell K_ATP_ channels are primarily regulated by glycolytic ATP^9^. The timing of oxidative phosphorylation, which is activated by ADP and reinforced by Ca^2+^ after membrane depolarization, does not align with the timing of K_ATP_ closure during pulsatile insulin secretion^10–14^ However, the activity of pyruvate kinase (PK), which converts ADP and PEP into ATP and pyruvate in the last step of glycolysis, correlates with the ATP/ADP rise preceding membrane depolarization and calcium influx^10–14^. It was recently discovered that a plasma membrane-associated pool of PK is sufficient to close K_ATP_ channels in human and mouse β cells^13^. Genetic studies of PK deficiency further suggest that mitochondrially-derived ATP buffers the cytosolic ATP/ADP ratio, but may not be the primary driver of ATP/ADP_pm_ or K_ATP_ closure^15^. It remains unclear whether compartmentalized regulation of K_ATP_ exists in α cells from humans or mice.

Compartmentation of glycolytic enzymes has been hypothesized to provide a highly efficient mechanism for localized regulation of the ATP/ADP ratio^16–19^. To date, the best evidence that glycolysis supports substrate channeling comes from protozoa – a special case where all nine glycolytic enzymes are contained within a membrane-bound organelle called the glycosome^20–22^. Under stress conditions, yeast are capable of assembling glycolytic enzymes into non-membrane bound granules termed G-bodies that accelerate glucose consumption^23,24^. In mammalian cells, lower glycolysis is known to regulate a variety of plasma membrane ion channels and pumps that exist in regions of high ATP consumption, including cardiac K_ATP_ channels^16,25–29^. However, it is not known whether upper glycolysis is also present and participates in K_ATP_ regulation, leaving the concept of a glycolytic metabolon elusive.

Here, we performed electrophysiological measurements of K_ATP_ activity in native α and β cell plasma membranes from both humans and mice. The excised patch clamp approach we employed not only identifies the plasma membrane localization of the endogenous enzymes, but also demonstrates their functional relationship to the activity of native K_ATP_ channels. Our experiments provide direct evidence that the enzymes of upper and lower glycolysis locally regulate K_ATP_ channels on the plasma membrane. Glucokinase (GK) and phosphofructokinase (PFK) provide ADP to activate K_ATP_ channels, while the enzymes of lower glycolysis utilize substrate channeling to produce ATP that closes K_ATP_. We further demonstrate the presence of two coupled reactions: 1) lactate dehydrogenase (LDH) that facilitates local NAD^+^/NADH recycling for GAPDH, as well as 2) the direct transfer of ADP generated by PFK to PK that ultimately closes K_ATP_ channels. These results identify a plasma membrane-associated glycolytic metabolon that locally signals via K_ATP_ channels to orchestrate α and β cell function.

## Results

### The plasma membrane-associated enzymes of lower glycolysis close K_ATP_ channels

We previously established that PK is present on the plasma membrane of human and mouse β cells where it raises ATP/ADP_pm_ sufficiently to close K_ATP_ channels^13,15^. Given that hormone secretion from both α and β cells relies on K_ATP_ channel regulation, we compared the effectiveness of substrate-driven K_ATP_ closure by PK in the two cell types. In dispersed mouse islets, the larger β cells were easily distinguished from α cells by size^7^, whereas anti-NTPDase3 coated magnetic beads were used to separate human α and β cells^30–32^.

As in our previous studies, we utilized the inside-out mode of excised patch clamp to study the effect of local ATP production by endogenous plasma membrane-associated PK on K_ATP_ channel activity (Fig. 1A). K_ATP_ channels were identified by closure upon the addition of bath solution containing 1 mM ATP, and reopening after switching to a high ADP solution containing 0.1 mM ATP and 0.5 mM ADP. In the presence of ADP, increasing concentrations of the PK substrate PEP (0.25, 1, and 5 mM) led to the dose-dependent closure of K_ATP_ channels in both mouse α and β cells (Fig. 1B, C). Channel closure occurred despite the continuous replacement of 0.5 mM ADP in the bath solution, which complexes with free Mg^2+^ to form MgADP that opens the K_ATP_ channel^2,3^. Thus, plasma membrane-compartmentalized PK must be sufficiently close to the K_ATP_ channel to locally deplete MgADP and increase ATP to close the channel. An alternative interpretation of this data is that PK-derived pyruvate is oxidized by residual patch-associated mitochondria that generate ATP and close K_ATP_ channels independently of glycolysis^33^. However, pyruvate and ADP were unable to induce K_ATP_ channel closure, and the presence of ATP synthase inhibitor (1 μg/mL oligomycin) did not affect K_ATP_ channel inhibition by PEP and ADP (Supplemental Figure S1). Consistently, knock-down of β cell PK activity abolished PEP-mediated K_ATP_ closure^15^. In most experiments, the PEP-dependent reduction in channel activity was accompanied by a lowered channel opening frequency without any change in event duration, both of which are shown for all treatments in Supplemental Table 1. Significant heterogeneity in the response to PEP treatment was observed between α and β cells (Fig. 1B, C).

**Figure 1.**
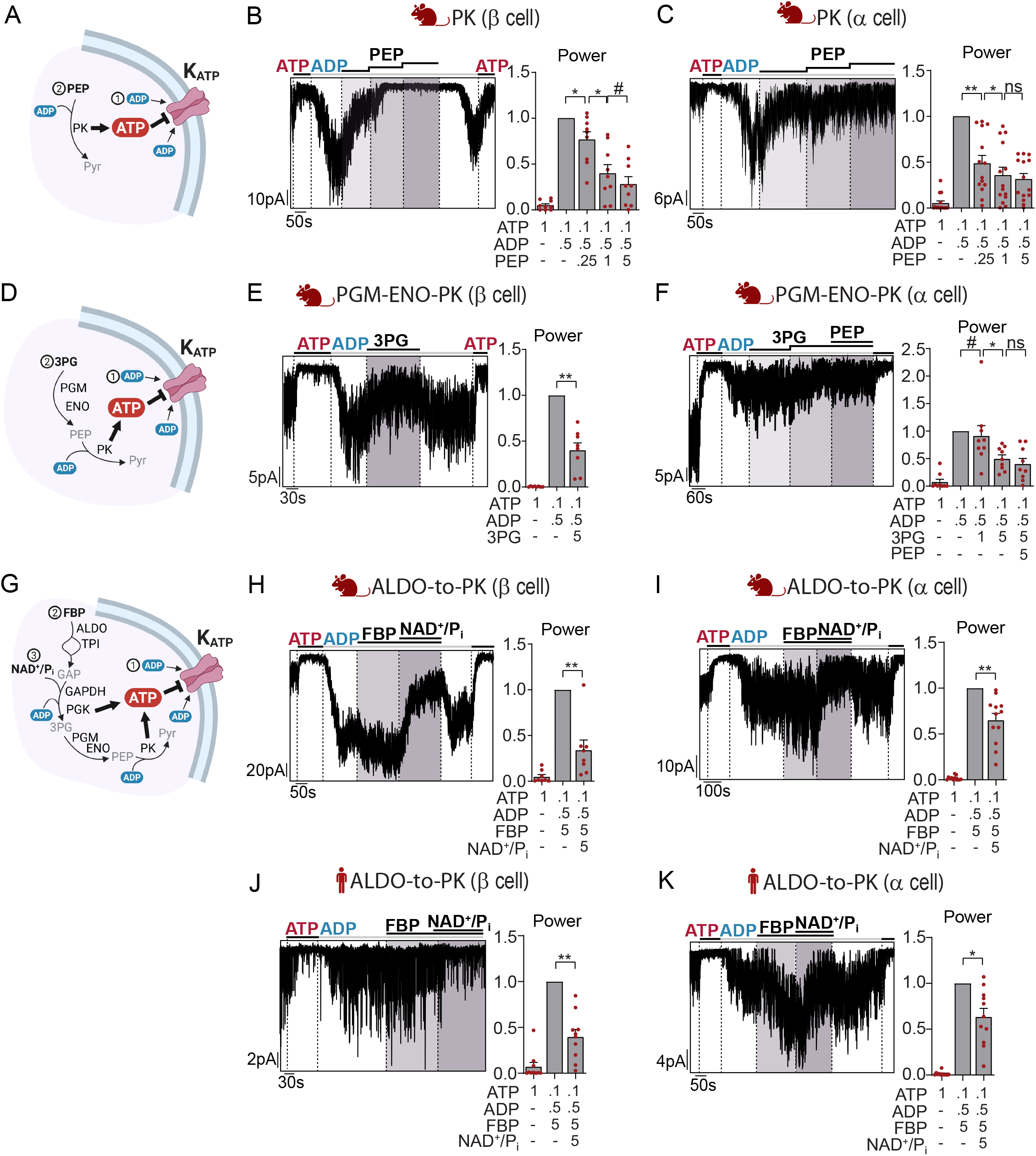
Lower glycolytic enzymes raise ATP/ADP to close α and β cell K_ATP_ channels. (A-C) PEP (0.25, 1, 5 mM) in the presence of high ADP (0.5 mM ADP, 0.1 mM ATP) led to the production of ATP by PK (A), thereby closing K_ATP_ channels in excised plasma membrane from mouse β cells (B) and mouse α cells (C). (D-F) In the high ADP condition, the chain reaction of PGM, ENO, and PK produced ATP and closed K_ATP_ channels (D) upon addition of 5 mM 3PG in mouse β cells (E) and 1 mM and 5 mM 3PG, followed by 5 mM PEP in mouse α cells (F). (G-K) Lower glycolytic activity (from ALDO to PK) utilized 5 mM FBP as the substrate for ALDO, 5 mM NAD^+^ and P_i_ (in the form of KH_2_PO_4_) as the substrate for GAPDH in high ADP condition to produce ATP (G), closing K_ATP_ channels in mouse β cells (H), mouse α cells (I), human β cells (J) and human α cells (K). Example traces show closure of K_ATP_ channel under 1 mM ATP, reopening after switching to high ADP condition, followed by addition of glycolytic substrates. Graphs quantify channel activity (power) normalized to (B-C, E-F, H-I) high ADP condition or (H-K) High ADP + 5 mM FBP condition. Data are from at least 3 mice or 3 human donors and shown as mean ± SEM. #p < 0.1, *p < 0.05, **p < 0.01, ***p < 0.001, ****p < 0.0001 by ratio paired student’s t test.

We next tested whether the two glycolytic enzymes upstream of PK, phosphoglycerate mutase (PGM) and enolase (ENO), could locally provide PEP for PK-mediated K_ATP_ closure on the plasma membrane (Fig. 1D). The application of the PGM substrate 3-phosphoglycerate (3PG, 5 mM) to a high ADP solution reduced K_ATP_ channel activity in mouse β cells (Fig. 1E) with similar efficacy to 5 mM PEP (Fig. 1B). In mouse α cells, we observed no effect of 1 mM 3PG but significant K_ATP_ closure with 5 mM 3PG (Fig. 1F). The further activation of PK with 5 mM PEP had no further effect. These findings indicate strong substrate channeling between the PGM-ENO-PK enzymes that provide substrate for PK to locally regulate K_ATP_ activity.

In cardiac myocytes, the glycolytic enzymes between aldolase (ALDO) and PK have been described to regulate K_ATP_ channels^26^. For comparison, we assessed K_ATP_ channel closure induced by the metabolism of the ALDO substrate FBP, which subsequently requires NAD^+^ and inorganic phosphate (P_i_) at the GAPDH step in order to produce ATP in the downstream PGK and PK reactions (Fig. 1G). Although the addition of 5 mM FBP alone led to variable changes in K_ATP_ channel activity, the addition of equimolar NAD^+^ and P_i_ (in the form of KH_2_PO_4_) led to strong inhibition of K_ATP_ channel activity in both mouse β and α cells (Fig. 1H, I), indicating that ATP production by lower glycolysis is coupled with K_ATP_ channel closure on the plasma membrane. Bath application of FBP to human β and α cell plasma membranes led to an activating effect on K_ATP_ channels. Here too, the further addition of NAD^+^ and P_i_ resulted in the inhibition of K_ATP_ channel activity in both β and α cells (Fig. 1J, K), indicating that the enzymes encompassing ALDO to PK are present on the plasma membrane where they function to regulate K_ATP_ channel activity at the ATP producing steps.

### The plasma membrane-associated enzymes of upper glycolysis open K_ATP_ channels

Multiple enzymes of upper glycolysis (hexokinase and glucose-6-phosphate isomerase) have been found by immunoprecipitation mass spectrometry to associate with the Kir6.2 subunits of the K_ATP_ channel^26^. However, it is not yet known whether upper glycolysis regulates K_ATP_ channel activity. We therefore assessed if ADP production by PFK could oppose K_ATP_ closure by a high ATP bath solution (1 mM ATP and 0.075 mM ADP) (Figure 2A). The application of fructose 6-phosphate (F6P, 10 mM) reversed K_ATP_ inhibition in mouse β and α cells (Fig. 2B, C), suggesting that MgADP reached a sufficient level in the K_ATP_ channel microcompartment to activate the nucleotide-binding domain of SUR1^34^. The K_ATP_ channel opening effect of F6P was found in ~40% of human β cells and ~60% of human α cells, but the level of α cell activation was approximately 3 times larger on average than β cells (Fig. 2E, F).

**Figure 2.**
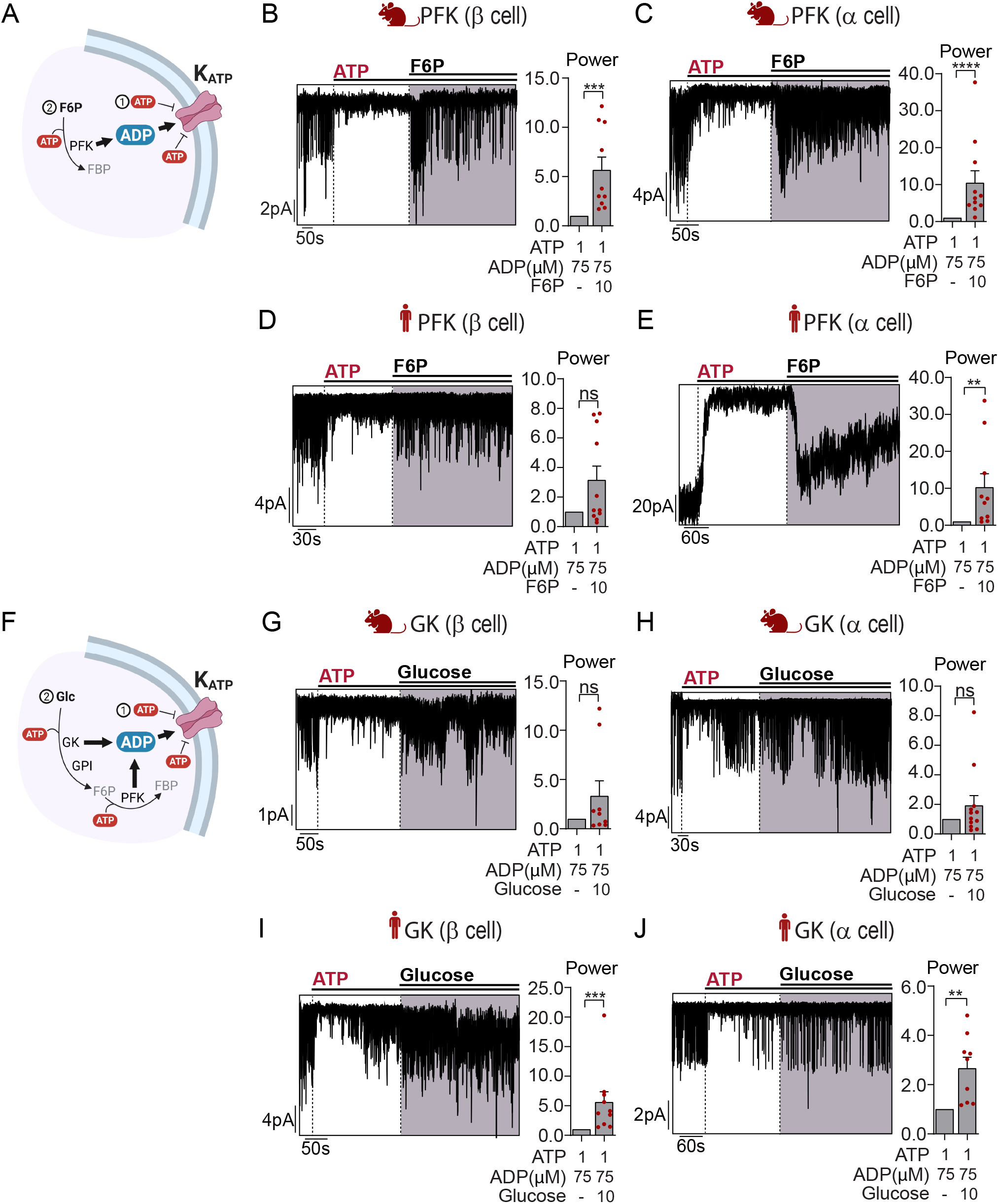
Upper glycolytic enzymes produce ADP that activates K_ATP_ channels. (A-E) In the high ATP condition (1 mM ATP, 0.075 mM ADP), addition of 10 mM F6P led to production of ADP by PFK (A), thereby opening K_ATP_ channels in mouse β cells (B), mouse α cells (C), human β cells (D) and human α cells (E). (F-J) Addition of 10 mM glucose in high ATP (F) produced little effect in mouse beta and α cells (G-H), but led to significant K_ATP_ activation through ADP production by GK in human β and α cells (I-J). Example recordings show K_ATP_ inhibition in the high ATP condition, followed by the addition of substrates for PFK and GK. Graphs quantify channel activity (power) normalized to high ATP condition. Data are from at least 3 mice or 3 human donors and shown as mean ± SEM. #p < 0.1, *p < 0.05, **p < 0.01, ***p < 0.001, ****p < 0.0001 by ratio paired student’s t test. Concentrations are in mM unless otherwise noted.

We next tested the coupling between glucokinase (GK)-derived ADP and the K_ATP_ channel (Figure 2F). In the presence of a high ATP/ADP ratio (1 mM ATP and 0.075 mM ADP), the addition of 10 mM glucose to mouse β and α cell plasma membranes led to significant K_ATP_ channel reopening in approximately 20% of the experimental replicates (Fig. 2G, H). In contrast, human islet cell membranes exhibited clear K_ATP_ opening in response to glucose, which was sufficient to overcome the high ATP/ADP ratio present in the bath solution (Fig. 2I, J). Together, these data demonstrate that the upper glycolytic enzymes are indeed present on the plasma membrane and capable of activating K_ATP_ channels.

### Although lactate activates K_ATP_, the presence of plasma membrane-associated LDH does not impede the ability of PK to close K_ATP_ channels

Lactate is a known metabolite activator of K_ATP_ channels^32,35–37^. Sarcolemmal K_ATP_ channels in inside-out excised patches are activated by 20 mM lactate, an effect that is reversed by the addition of equimolar NAD^+^ to remove lactate by the LDH reaction^36^. Because the circulatory level of lactate does not rise above 10 mM except during intense exercise^38^, we tested whether a lower concentration of lactate (2 mM) could produce the same effect (Supplemental Figure S2A). Lactate application to mouse β cell plasma membranes in the high ATP condition (1 mM ATP and 0.075 mM ADP) had a heterogeneous effect, increasing K_ATP_ activity significantly in 3 of 11 experiments (Supplemental Figure S2B). However, in mouse α cell membranes, we observed significant activation of K_ATP_ channels at 2 mM lactate (Supplemental Figure S2C). The addition of excess NAD^+^ (5 mM) abolished the activating effect of lactate, indicating the presence of LDH on α cell plasma membrane. In excised patches from human islet cells, we observed dose-dependent opening of K_ATP_ channels after the addition of 2 and 5 mM lactate in both α and β cells, however the addition of 5 mM NAD^+^ had no further effect (Supplemental Figure S2D, E). In summary, exogenous lactate has a strong K_ATP_ channel opening effect in both mouse and human α and β cells, and we confirmed plasma membrane LDH activity in mouse α cells (and see next section for the effect of LDH in β cells).

Given the ability of lactate to open K_ATP_ channels and hyperpolarize mouse α cells that express MCT1 pyruvate/lactate transporters^32^, we next tested whether exogenous lactate at concentrations mimicking circulatory levels is potent enough to oppose PK-mediated K_ATP_ channel closure in mouse α cell membranes. The application of 2 mM lactate was able to reopen K_ATP_ channels inhibited by 2 mM PEP and 0.5 mM ADP (Supplemental Figure S3A, B). However, it is unlikely that K_ATP_ channels are regulated by LDH-derived lactate, since the addition of NADH (2 mM) was unable to reverse K_ATP_ closure by PK (Supplemental Figure S3C, D). Pyruvate (2 mM) also activated K_ATP_ channels in a subset of α cell plasma membranes, and again, equimolar NADH had no additional effect (Supplemental Figure S3E, F). These data argue that, despite the presence of LDH activity in the K_ATP_ microcompartment, LDH-mediated lactate production is insufficient to overcome K_ATP_ channel closure by PK even in mouse α cells with high LDH activity.

### LDH and GAPDH facilitate a plasma membrane-associated NAD^+^/NADH cycle that supports K_ATP_ channel closure by PK

One of the main functions of the LDH reaction is to regenerate NAD^+^, which is necessary for the continuous oxidation of glucose at the GAPDH step. Given the presence of LDH on the plasma membrane, we hypothesized the formation of a glycolytic metabolon that not only facilitates the channeling of metabolites between enzymes, but also allows NAD^+^/NADH cycling within the K_ATP_ channel micro-compartment to maintain redox balance. In order to test whether LDH generates sufficient NAD^+^ to support ATP generation by lower glycolysis, we monitored K_ATP_ activity in response to FBP metabolism. In this experiment, pyruvate and NADH (5 mM each) were provided as substrates for LDH, and 5 mM P_i_ was provided for the GAPDH reaction, such that the only source of NAD^+^ is the LDH reaction (Figure 3A). The observed reduction in K_ATP_ channel activity in both human and mouse α and β cells (Fig. 3B-E), which we interpret to be a result of FBP metabolism by lower glycolysis leading to an increase in ATP/ADP_pm_, was very similar to when FBP, NAD^+^, P_i_ and ADP were provided directly (Fig. 1H-K). These findings demonstrate that plasma membrane-associated LDH and GAPDH facilitate a local NAD^+^/NADH redox cycle within the K_ATP_ channel micro-compartment.

**Figure 3.**
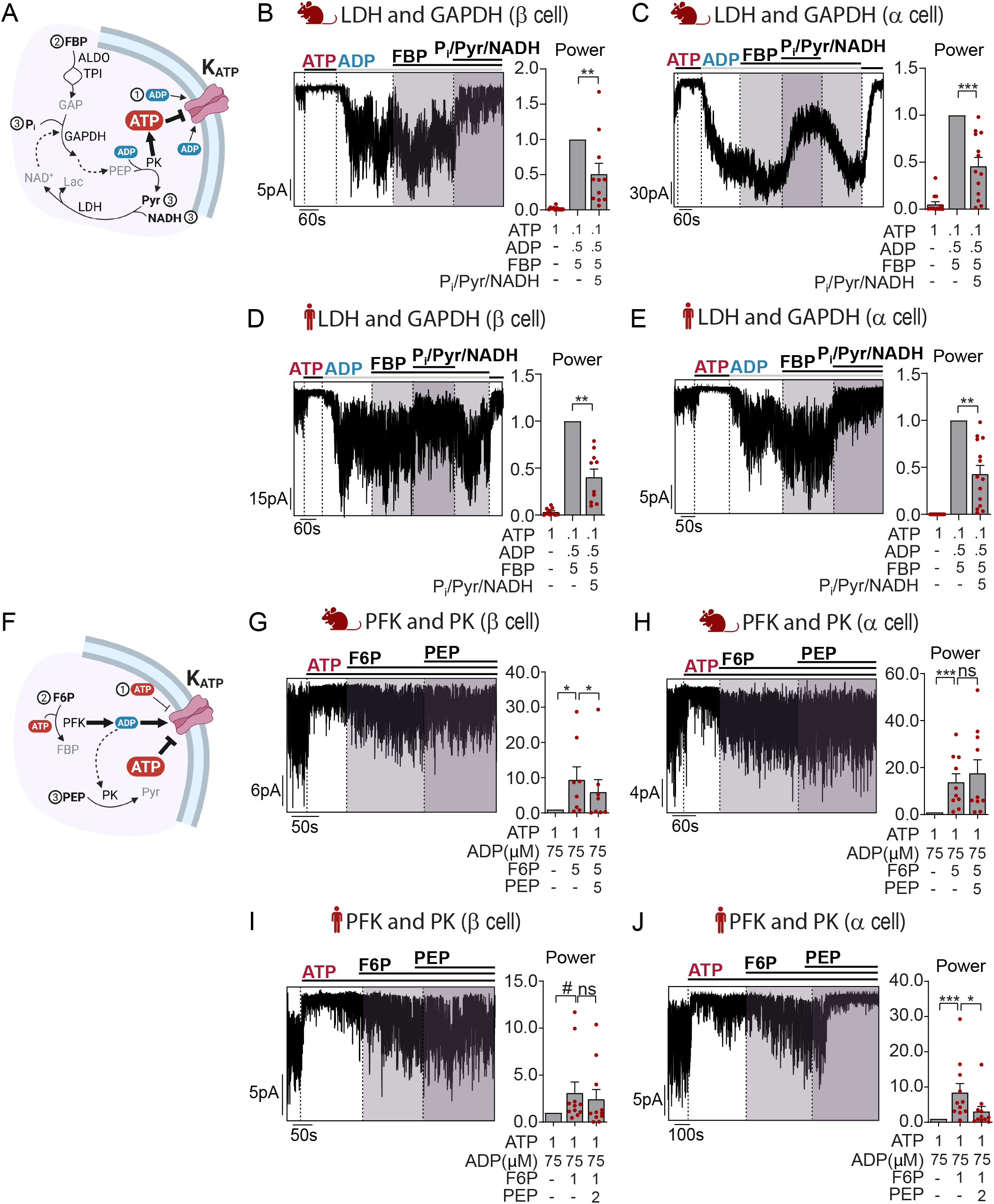
The ADP produced by PFK is used directly by PK, and NAD^+^ produced by plasma membrane-associated LDH is used directly by GAPDH to sustain ATP production. (A-E) Addition of 5 mM pyruvate (Pyr) and NADH allows LDH reaction to provide NAD^+^ at the GAPDH step to metabolize 5 mM FBP (substrate for ALDO) to ATP that closes K_ATP_ channels in mouse and human β and α cells. (F-J) Addition of F6P (5 mM in mice, 1 mM in humans) under high ATP condition (1 mM ATP, 0.075 mM ADP) led to activation of K_ATP_ channels; the subsequent addition of PEP (5 mM in mice, 2 mM in humans) did not significantly alter K_ATP_ activity in mouse β (G-H) or human (I) β cells, but led to K_ATP_ inhibition through PK-generated ATP in human α cells (J). Example traces show K_ATP_ inhibition in 1 mM ATP and reopening in high ADP condition (0.5 mM ADP, 0.1 mM ATP) (B-E) K_ATP_ inhibition in high ATP condition (G-J), followed by the addition of test substrates. Graphs quantify channel activity (power) normalized to high ADP + 5 mM FBP condition (B-E) or high ATP condition (G-J). Data are from at least 3 human donors and shown as mean ± SEM. #p < 0.1, *p < 0.05, **p < 0.01, ***p < 0.001, ****p < 0.0001 by ratio paired student’s t test. Concentrations are in mM unless otherwise noted.

### The ADP produced by PFK is utilized directly by PK to close K_ATP_ channels

As ATP/ADP_pm_ rises and flux through PK slows, there is a greater need to provide ADP to the PK reaction to sustain glucose oxidation. Because the GK and PFK reactions in upper glycolysis consume ATP, we hypothesized that one or both enzymes could act as a direct source of ADP substrate for PK (Figure 3F). To test this, we first drove ADP production by the PFK reaction using F6P (5 mM in mice, 1 mM in humans), resulting in the opening of K_ATP_ channels under a high ATP/ADP ratio (1 mM ATP and 0.075 mM ADP) in the bath solution (Fig. 3G-J). In these experiments, the F6P concentration was much lower than in Fig. 2B-E, reflecting a more physiological level. While we did not observe coupling between PFK and PK in mouse α cells or human β cells, the subsequent application of PEP (5 mM in mouse, 2 mM in human) closed K_ATP_ channels in mouse β cells and human α cells, demonstrating that ADP was converted to ATP by the PK reaction (Fig. 3G, H, I and J).

## Discussion

Our findings show that a glycolytic metabolon—reflecting a functional signaling machinery but not necessarily a unitary complex—is localized to the plasma membrane of α and β cells where it locally controls the activity of K_ATP_ channels. We detected K_ATP_ channel regulation by the rate-controlling enzymes of upper and lower glycolysis as well as LDH. Efficient substrate channeling of FBP was observed between the enzymes of lower glycolysis, in addition to the direct transfer of ADP between the PFK and PK reactions. Furthermore, we elucidated a plasma membrane NAD^+^/NADH cycle mediated by NAD^+^ channeling between LDH and GAPDH. Despite small differences, this strategy of compartmentalized K_ATP_ channel regulation is mostly conserved between humans and mice. In all treatments, we routinely observed variability between cells, which provides a plausible explanation for the well-documented metabolic heterogeneity within populations of α and β cells^39–42^.

We found that two enzymes in upper glycolysis, GK and PFK, locally activate K_ATP_ channels by raising the level of ADP in sub-plasma membrane space. ADP then serves as a direct substrate for ATP generation by PK. While PGK is also present and provides a source of ATP, this near-equilibrium enzyme lacks metabolic control^43^. Although glycolysis consumes 2 and generates 4 ATP, this does not necessarily imply that the enzymes of upper glycolysis are irrelevant to K_ATP_ channel regulation. In β cells, the coupling efficiency of upper and lower glycolysis can be modified by the siphoning of substrates. In individuals with type 2 diabetes, after GK produces ADP, glucose-6-phosphate is shuttled towards glycogen synthesis and thus decreases glucose-stimulated insulin secretion through reduced ATP output^44^. Similarly, glycerol-3-phosphate phosphatase has been shown to operate a glycerol shunt to curtail hyperinsulinemia by removing dihydroxyacetone phosphate from glycolysis, which can reduce ATP generation up to 50%^45,46^. Further work is necessary to identify whether these metabolic shunts are present in the plasma membrane or whether they occur in another subcellular compartment.

In addition to glycolysis, LDH activity was found to be present in the K_ATP_ microcompartment and efficiently drive NAD^+^ production to support glycolysis at the GAPDH step in both human α and β cell membranes. In support of previously published work^32,35–37^, we found that lactate opens K_ATP_ channels and opposes PK-mediated K_ATP_ closures in mouse α cells. Since lactate can reach millimolar levels in the blood, and α-cells express a high level of MCT pyruvate/lactate transporters (*Slc16a1*)^32^, lactate regulation of K_ATP_ channels is expected to occur in vivo. However, lactate derived from glycolytic pyruvate had no impact on K_ATP_ channels in our experiments. In β cells, *Ldha* is considered a “disallowed gene”^47–49^. However, human β cells express two isoforms of LDH (A/B)^50^, and ectopic expression of MCT1 is sufficient for lactate-induced insulin secretion that mediates exercise-induced hyperinsulinemic hypoglycemia^51^. Our results further demonstrate that lactate production provides NAD^+^ to support K_ATP_ channel closure. We previously reported glucose-dependent lactate production and lactate oscillations in mouse islet β cells^52^, demonstrating the physiological relevance of these findings.

Pancreatic endocrine cells, like cardiac myocytes, may share K_ATP_ as the anchor point for lower glycolysis, however a diversity of membrane scaffolds have been uncovered^53–56^. Disrupting this association would help the islet field determine whether stimulus-secretion coupling remains efficient without glycolytic compartmentation. In addition, we do not yet know the relative importance of glycolytic versus mitochondrial K_ATP_ regulation in islet cells from individuals with diabetes. While we attempted to utilize frozen islet samples from human donors with type 2 diabetes, the thawed tissues were not viable for excised patch-clamp experiments after shipping. Nonetheless, our prior work suggests the potential of small molecule PK activators for the treatment of diabetes based on the local coupling between PK and K_ATP_ channels^13,57^. Although current research in pancreatic hormone secretion and diabetes largely overlooks the distinction between metabolism in the bulk cytosol and micro-compartments, partly due to a lack of suitable tools, our results highlight the need for further work on metabolic compartmentation that could become the basis for novel therapeutic strategies.

## Supporting information

List of human donors

Excised patch clamp frequency and duration statistics

## Figure legends

**Supplementary Figure S1.**
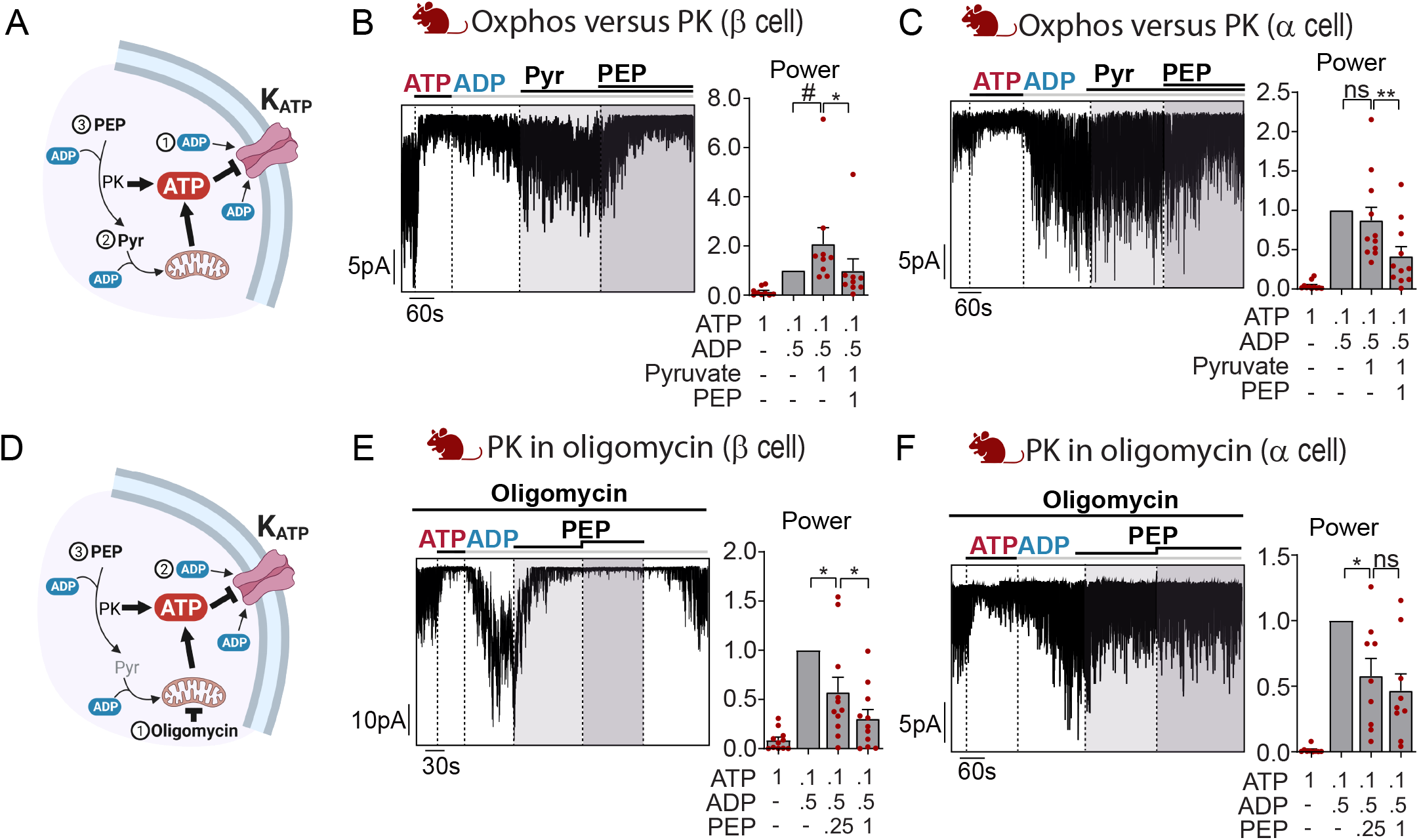
PEP-mediated K_ATP_ closure is independent of mitochondrial oxidative phosphorylation. (A-C) Addition of 1 mM pyruvate (Pyr) under high ADP condition (0.5 mM ADP, 0.1 mM ATP) was not sufficient to induce K_ATP_ channel closure; but subsequent addition of 1 mM PEP allowed for PK-mediated ATP generation that inhibits K_ATP_ channels in mouse β and α cells. (D-F) K_ATP_ closure induced by PEP (0.25, 1 mM) under high ADP condition (0.5 mM ADP, 0.1 mM ATP) was not affected by the presence of the ATP synthase inhibitor (oligomycin, 1 μg/mL) in both mouse β and α cells. Example traces show closure of K_ATP_ channel under 1 mM ATP, reopening after switching to high ADP condition, followed by addition of glycolytic substrates. Graphs quantify channel activity (power) normalized to high ADP condition. Data are from at least 3 mice and shown as mean ± SEM. #p < 0.1, *p < 0.05, **p < 0.01, ***p < 0.001, ****p < 0.0001 by ratio paired student’s t test. Concentrations are in mM unless otherwise noted.

**Supplementary Figure S2.**
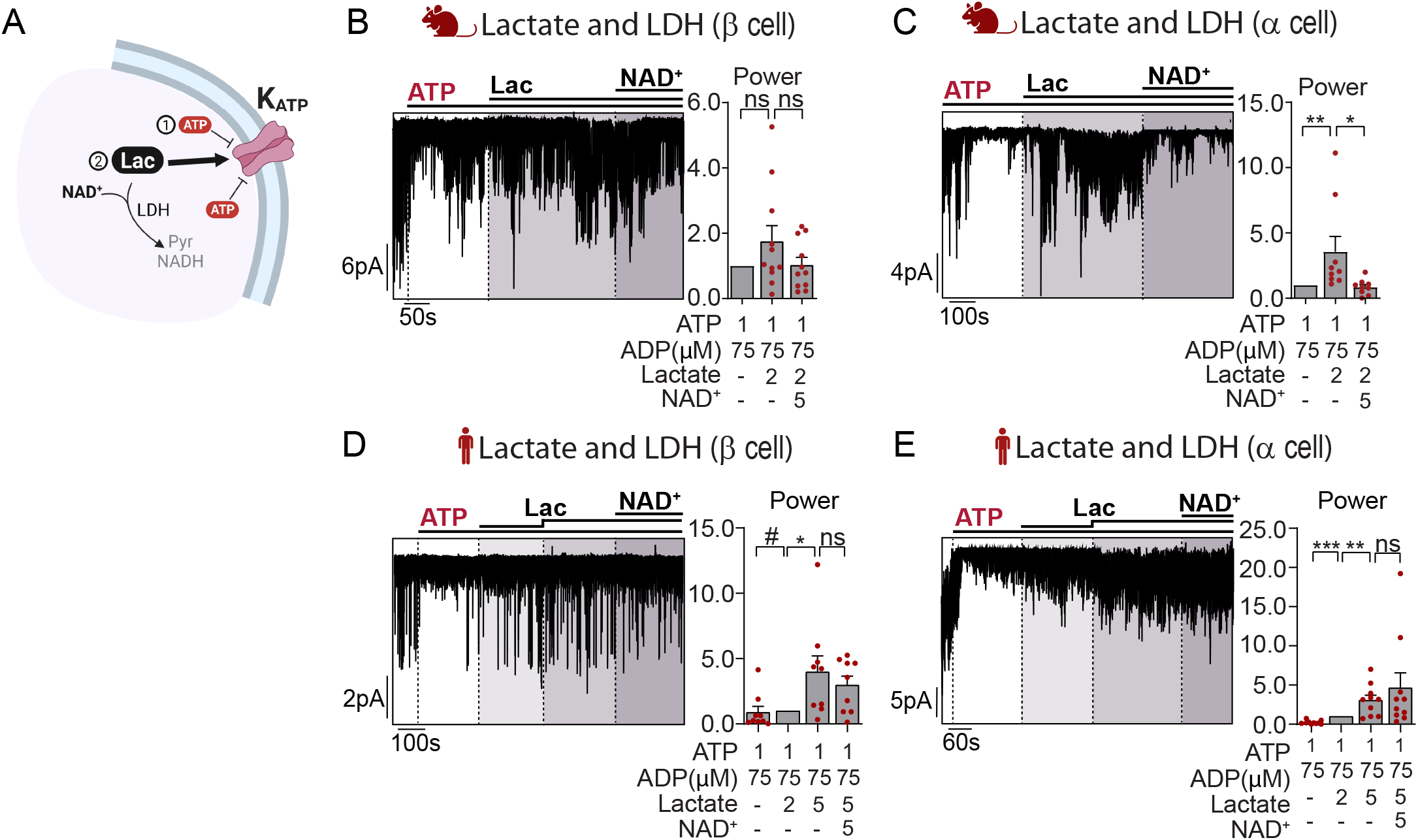
Exogenous lactate opens K_ATP_ channels. (A) Addition of lactate (2 mM) opens K_ATP_ channels in high ATP condition (1 mM ATP, 0.075 mM ADP) followed by excess NAD^+^ (5mM) for the LDH reaction did not lead to changes in K_ATP_ channel activity in (B) mouse β cells, but had a significant activating effect with lactate and in (C) mouse α cells. (D-E) Lactate (2, 5 mM) opens K_ATP_ channels in human β and α cells but equimolar NAD^+^ (5mM) had no further effect. Example traces show K_ATP_ closure under high ATP condition, followed by application of lactate and NAD^+^. Graphs quantify channel activity (power) normalized to (B-C) high ATP condition or (D-E) 2 mM lactate condition. Data are from at least 3 mice or 3 human donors and shown as mean ± SEM. #p < 0.1, *p < 0.05, **p < 0.01, ***p < 0.001, ****p < 0.0001 by ratio paired student’s t test. Concentrations are in mM unless otherwise noted.

**Supplementary Figure S3.**
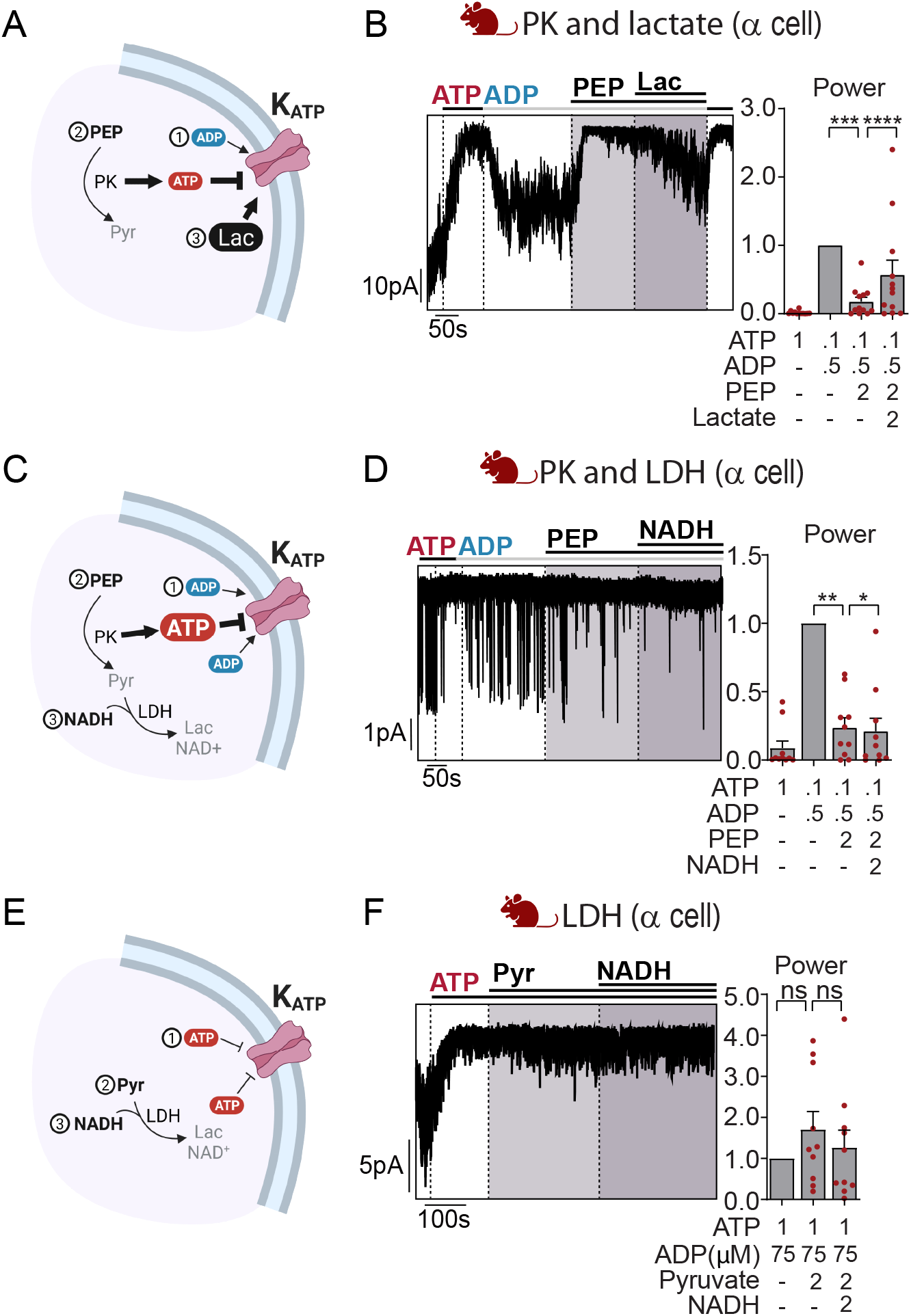
Exogenous lactate opposes PEP-mediated closure, however endogenous LDH activity is insufficient to activate K_ATP_ channels. (A-B) In mouse α cells, addition of 2 mM PEP in high ADP condition (0.5 mM ADP, 0.1 mM ATP) led to K_ATP_ channel closure through the PK reaction; subsequent addition of equimolar lactate (2 mM) reopens K_ATP_ channels. (C-D) Following K_ATP_ closure mediated by 2 mM PEP in high ADP condition, further addition of equimolar NADH (2 mM) to activate the LDH reaction failed to reopen K_ATP_ channels in mouse α cells. (E-F) In the high ATP condition (1 mM ATP, 0.075 mM ADP), addition of direct substrates for LDH to produce lactate (2 mM pyruvate and NADH) failed to produce significant K_ATP_ channel opening in mouse α cells. Example traces show (B and D) K_ATP_ channel closure in 1 mM ATP and subsequent reopening with high ADP condition followed by application of substrates; or (F) K_ATP_ closure in high ATP condition followed by application of substrates. Graphs quantify channel activity (power) normalized to (B and D) high ADP condition or (F) high ATP condition. Data are from at least 3 mice and shown as mean ± SEM. #p < 0.1, *p < 0.05, **p < 0.01, ***p < 0.001, ****p < 0.0001 by ratio paired student’s t test. Concentrations are in mM unless otherwise noted.

**Supplementary Table 1. Normalized average of event frequency and single event duration** Normalized average values are listed as mean ± SEM and in the same order as their respective power quantification.

## Acknowledgments

The Merrins laboratory gratefully acknowledges support from the NIH/NIDDK (R01DK113103 and R01DK127637). This work was supported in part by the United States Department of Veterans Affairs Biomedical Laboratory Research and Development Service (I01B005113). We would like to thank Dudley Lamming for contributing support for EP from R01AG062328. This work utilized facilities and resources from the William S. Middleton Memorial Veterans Hospital and does not represent the views of the Department of Veterans Affairs or the United States Government. Graphics were created using BioRender.com. We thank the Human Organ Procurement and Exchange (HOPE) program and Trillium Gift of Life Network (TGLN) for their work in procuring human donor pancreas for research, and James Lyon, Dr. Jocelyn Manning Fox, and Dr. Patrick MacDonald (Alberta) for their efforts in human islet isolation. We especially thank the organ donors and their families for their kind gifts in support of diabetes research.

## Author Contributions

M.J.M. conceived the study and wrote the paper with T.H. T.H. performed the main body of experiments with assistance from E.P., D.B.D., and M.J.M. All authors interpreted the data and edited the manuscript.

## STAR methods

### Resource Availability

#### Lead Contact

Further information and requests for resources and reagents should be directed to and will be fulfilled by the Lead Contact, Matthew Merrins.

#### Materials Availability

This study did not generate new unique reagents.

#### Data and Code Availability

This study did not generate new code.

### Experimental Model and Subject Details

#### Mouse Islet Preparations

C57BL/6J male mice were ordered from the Jackson Laboratory and housed and used as previously described. The mice are sacrificed by cervical dislocation and islet isolations were carried out as previously described (Gregg et al., 2016). Islets were cultured in RPMI-1640 supplemented with 10% (v/v) fetal bovine serum (ThermoFisher A31605), 10,000 units/mL penicillin and 10,000 mg/mL streptomycin (Fisher Scientific). All procedures involving animals were approved by the Institutional Animal Care and Use Committees of the University of Wisconsin-Madison and the William S. Middleton Memorial Veterans Hospital, and followed the NIH Guide for the Care and Use of Laboratory Animals (8th ed. The National Academies Press. 2011.).

#### Human Islet Preparations

Human islets from normal donors were obtained from the Alberta Diabetes Institute IsletCore and the Integrated Islet Distribution Program. The age, sex, body mass index, and %HbA1c of each donor are provided in Supplementary Table S2. Human islets were cultured in glutamine-free CMRL supplemented with 10 mM niacinamide and 16.7 μM zinc sulfate (Sigma), 1% ITS supplement (Corning), 5 mM sodium pyruvate, 1% Glutamax, 25 mM HEPES (American Bio), 10% heat-inactivated FBS and antibiotics (10,000 units/mL penicillin and 10 mg/mL streptomycin).

### Method Details

#### Islet dispersion

Islets were washed briefly with VERSENE solution (ThermoFisher 15040066), and dissociated using TrypLE Select Enzyme solution (ThermoFisher 12563011) at 37°C, with occasional trituration. For human islets, the dissociated cells were purified into β cell and non-β cell fractions as described previously using mouse anti-human NTPDase3 antibodies (Ectonucleotidases-ab) and CELLection Pan Mouse IgG kit (ThermoFisher 11531D)^32^. Cells were plated on sterilized glass shards and cultured overnight at 37°C before experiments. For mouse cells, culture media was RPMI-1640 supplemented with 10% (v/v) fetal bovine serum (ThermoFisher A31605), 10,000 units/mL penicillin and 10,000 mg/mL streptomycin (Fisher Scientific). For human cells, the base media was glucose-free RPMI-1640 supplemented with 8 mM glucose.

#### Electrophysiology

A HEKA Instruments EPC10 patch-clamp amplifier was used for the registration of current. Intracellular recording electrodes made of borosilicate glass (Harvard Apparatus, Holliston, MA) were polished by a microforge (Narishige MF-830) to the final tip resistance of 4-10 MOhm. The pipette solution contained (in mM): 10 sucrose, 130 KCl, 2 CaCl_2_, 10 EGTA, 20 HEPES, pH 7.2, adjusted with KOH. Recordings were made at room temperature. Briefly, gigaseals (>2.5 GOhm) were established in extracellular bath solution (in mM): 140 NaCl, 5 KCl, 1.2 MgCl_2_, 2.5 CaCl_2_, 0.5 glucose, 10 HEPES, pH 7.4, adjusted with NaOH and held at −50 mV before excision into inside-out configuration. Equilibrium solutions with K^+^ as the charge carrier were used for recording. The recording bath solution (control) contained (in mM): 130 KCl, 2 CaCl_2_, 10 EGTA, 1.0 free Mg^2+^, 10 sucrose, 20 HEPES, pH 7.2 with KOH. ATP, ADP, and metabolites were added to the control solution and pH values were adjusted with KOH to 7.2. Data was filtered online at 1 kHz with a Bessel filter and analyzed offline using Clampfit 10 software (Axon Instruments). A 60-second window at the end of each condition was used to analyze K_ATP_ activity in terms of power, frequency, and duration, all of which were normalized to control conditions (typically the high ATP or high ADP condition) as indicated in the text. Akin to area under the curve, power is calculated as the product of the channel amplitude and single event duration, summed over all channels. Consequently, power is a proxy for the total number of ions passing through the channel, as is referred to as channel activity in the text. Frequency reflects the number of opening events throughout the analysis window while event duration measures the average length of time before closure occurs. The latter two parameters, which are reported in the supplemental data, only account for events that at least open or close during the window of analysis (i.e. they do not include always-active events) and thus are underestimations for conditions that have many always-opened channels.

